# Molecular profiling implicates immune-driven adaptive homeostasis in maintaining intraocular pressure with age

**DOI:** 10.64898/2026.08.19.745748

**Authors:** Guorong Li, Nicholas Tolman, Abdul Hannan, Joshua B. Ramkissoon, Justin Thomas Mathew, Karina Polanco, Kristin Perkumas, Maria Fernanda Suarez, Serena J. Cui, Roshnni Rajkumar, Christa Montgomery, Krishnakumar Kizhatil, W. Daniel Stamer, Simon W.M. John, Revathi Balasubramanian

## Abstract

Aging and ocular hypertension are primary risk factors for glaucoma - a major cause of blindness worldwide. Ocular hypertension results from dysfunction in the outflow tissues - the trabecular meshwork (TM) and Schlemm’s canal (SC), which are key regulators of intraocular pressure (IOP) homeostasis. Despite lifelong multifaceted stress to the outflow pathway, ocular hypertension sufficient to develop glaucoma occurs in only a minority of individuals. The mechanisms that preserve physiological IOP with aging in most individuals remain poorly understood. Using single-cell RNA sequencing and protein validation in mouse outflow tissues, we found that aging is accompanied by subtype-specific changes in TM cells, including altered extracellular matrix maintenance, reduced trophic signaling to SC, and increased profibrotic signaling. Coincident with these changes, we observed age-dependent immune remodeling, marked by increased macrophage abundance in outflow tissues. Our model predicts that macrophages maintain SC homeostasis via VEGFA signaling - normally supplied by the TM and required for normal SC function, but impaired with age. This pattern supports a paradigm in which adaptive immune remodeling contributes to preserving IOP homeostasis despite progressive aging-associated cellular dysfunction. More broadly, we identify the TM/SC outflow pathway as a model for understanding how aging tissues preserve physiological function through coordinated structural, trophic, and adaptive immune remodeling.

## Introduction

Age and elevated intraocular pressure (IOP) are primary risk factors for glaucoma - a leading cause of irreversible blindness^1–4^. In healthy people, IOP is stable within a couple of millimeters of mercury over a lifetime due to continual adjustments in outflow resistance by the drainage tissues - the trabecular meshwork (TM) and Schlemm’s canal (SC)^5–9^. However, IOP in people with glaucoma often increases with age resulting from increased resistance due to dysfunctional outflow homeostasis.^10–13^.

Like humans, IOP in healthy mice is relatively stable with age^14^. Outflow facility is also largely stable with age despite senescence-related tissue remodeling and deterioration in mouse drainage tissues^14–16^. Specifically, the TM and SC of elderly mice display decreased cellularity, increased pigment accumulation, increased cellular senescence and increased stiffness, similar to changes observed in human eyes with age and glaucoma^14,17–19^. Mice are also like humans in their anatomy, physiology and pharmacology of drainage tissues^20–26^. Such parallels with humans, in addition to their short lifespan, make mice a robust model to study age-related changes in outflow tissues. This is particularly important for outflow tissues where multiple unique cell types (mesenchymal, endothelial and immune cells) work together to maintain IOP over a lifetime^27–32^. To understand how the TM and SC (mechanically, metabolically and oxidatively stressed, immune-interacting tissues) preserve function during aging, we performed single-cell RNA sequencing of drainage tissues in three age groups of mice. We focused on age-related changes in the TM and SC, identifying key alterations in extracellular matrix remodeling, structural rigidity, cell death, and loss of essential trophic support. These findings were validated at the protein level using complementary methodologies. We propose a model in which macrophages compensate for the loss of trophic support to help maintain physiological homeostasis.

## Results

### A single-cell transcriptomic atlas of TM and SC from young, middle-aged, and old mice

To characterize the transcriptomic landscape of the outflow tissues across their lifespan, we performed large-scale single-cell (scRNA-seq) of limbal strips enriched for the SC and TM (Figure 1A). We profiled approximately 95,000 cells derived from limbal strips spanning three age groups: young (3 months), middle-aged (12 months), and old (22– 24 months). Isolation of cells and sequencing runs were conducted across two independent sites (Duke University and Columbia University) and successfully integrated.

**Figure 1:**
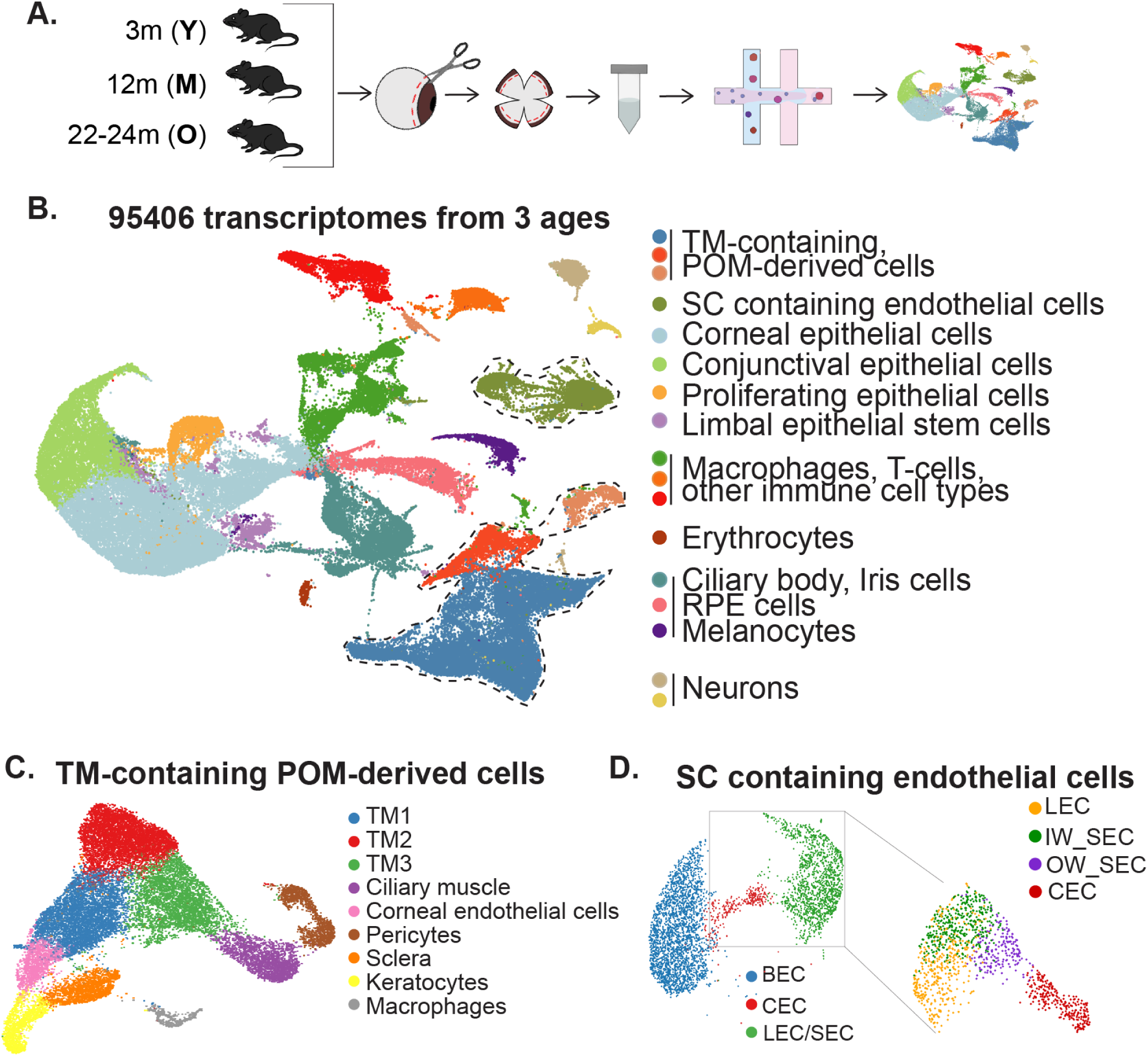
Large scale scRNA-seq of limbal strip tissues reveal distinct cell types. **(A)** A schematic representation of the experimental workflow used for scRNA-seq. Limbal strip derived cells from mice in 3 age cohorts (young, middle and old) were isolated, enriched for drainage tissues, and sequenced. **(B)** UMAP clusters of all major anterior segment cell types from scRNA-seq from 95,000 cells across all 3 age groups. **(C)** Iterative subclustering of the TM-containing POM-derived cell clusters reveals 3 distinct subtypes of TM cells (TM1, TM2, and TM3). **(D)** Iterative subclustering of SC-containing endothelial cell clusters reveals distinct subtypes of SC endothelial cells corresponding with the inner and outer wall, as well as other endothelial cell populations (blood, lymphatic, and circulating ECs).

Dimensionality reduction and visualization via Uniform Manifold Approximation and Projection (UMAP) enabled the systematic identification of major anterior segment cell types within the integrated dataset (Figure 1B, S1A). Attention was directed toward two biologically critical populations: the cluster enriched for periocular mesenchyme (POM)- derived TM cells, and the endothelial cell cluster containing SC cells. To refine the cellular identities within these compartments and delineate transcriptionally distinct subpopulations, we performed iterative sub-clustering. We resolved three transcriptionally distinct TM subtypes (TM1, TM2, and TM3), as well as discrete SC endothelial cell (SEC) subpopulations corresponding to the inner wall (IW) and outer wall (OW) of SC (Figure 1C, D, S2A). Together, these analyses provide a high-resolution cellular taxonomy of the outflow tissues and establish a framework for interrogating age-associated transcriptomic changes within SC and TM cell populations.

### Aging drives subtype-dependent gene expression changes across TM populations

To interrogate the molecular mechanisms underlying age-related dysfunction in the outflow pathway, we performed gene ontology (GO) pathway enrichment analysis and linear differential gene expression analysis across the three transcriptionally distinct TM subtypes (TM1, TM2, and TM3) identified in our dataset (Figure S3A). These subtypes are consistent with our previously published characterization^29^ of mouse TM subtypes (defined by both anatomical location and differential marker expression). We determined whether these subtype populations undergo distinct transcriptomic changes with age.

In TM1 cells, we observed a significant age-dependent downregulation of pathways associated with extracellular matrix (ECM) organization (Figure 2A). Focused analysis of collagen gene expression revealed broad transcriptional decreases across collagen family members implicated in both structural reinforcement of the ECM and intercellular signaling, suggesting a progressive deterioration of ECM architecture within the TM with age (Figure 2B).

**Figure 2:**
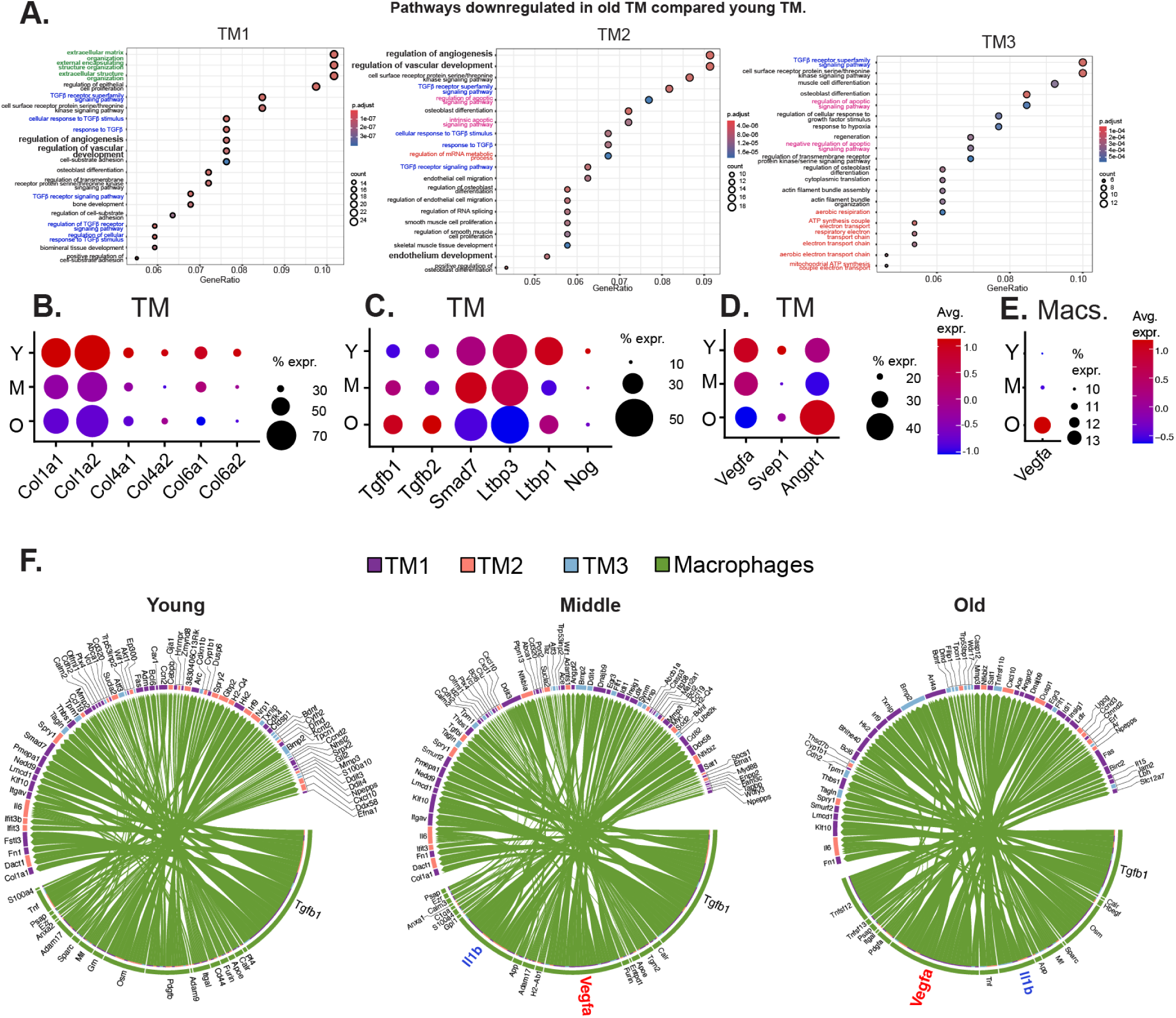
**(A)** GO pathway enrichment analysis across 3 TM subtypes. **(B)** Expression levels and percent cells expressing collagen genes across age groups in TM cells. **(C)** Expression levels and percent cells expressing genes involved in TGF-beta signaling pathway across age groups in TM cells. **(D)** Expression levels and percent cells expressing gene involved in angiogenic and vascular maintenance factors across ages in TM cells. **E)** Expression levels and percent cells expressing Vegfa in macrophages (Macs.). **(F)** Circos plot showing the top predicted signaling interactions between TM1, TM2, TM3 (receivers) and macrophage cells (senders) across age groups.

We further identified a downregulation of TGF-beta signaling pathway components across all three TM subtypes with age (Figure 2A). Given the complexity of TGF-beta signaling (encompassing several genes distributed across multiple regulatory nodes), we focused our analysis on a curated set of pathway genes. Specifically, we examined canonical ligand genes (*Tgfb1, Tgfb2*) and negative regulators of the pathway (*Smad7, Ltbp1, Ltbp3,* and *Nog*). Paradoxically, despite the overall downregulation of TGF-beta pathway gene sets by GO analysis, we observed an age-dependent increase in *Tgfb1* and *Tgfb2* expression, accompanied by a concurrent decrease in inhibitory regulators (Figure 2C). This net shift toward a pro-TGF-beta signaling environment is consistent with recent reports of elevated TGF-beta levels in glaucomatous ocular tissues, implicating dysregulated TGF-beta activity as a conserved feature of age-related TM pathology.

Additionally, GO analysis revealed an age-associated increase in apoptosis and reduction in paracrine signaling pathways governing vascular development and angiogenesis across TM subtypes, suggestive of declining trophic support to the adjacent SC with age (Figure 2A). Previous studies have shown that the TM functions as a supportive niche, secreting trophic factors necessary for normal SC function^30,33,34^. We examined the expression of key angiogenic and vascular maintenance factors in the TM, including vascular endothelial growth factors (Vegfs), angiopoietins (Angpts), and *Svep1*. We identified significant age-dependent decreases in *Vegfa* and the glaucoma-linked extracellular matrix protein gene *Svep1*^34–36^, which supports Schlemm’s canal.

Overall, however, *Angpt1* expression increased in old TM relative to young; however this trend was non-linear, with an apparent decrease at middle age followed by an elevation in expression in old age (Figure 2D).

Our previous study^14^ reported that neither IOP nor outflow facility is significantly altered in aged mice, raising the question of whether compensatory mechanisms exist to sustain SC function in the context of declining TM-derived trophic support. One possibility is that the downregulation of *Vegfa* may trigger a delayed *Angpt1* upregulation as a compensatory response. Another hypothesis is that non-TM cell populations within the outflow niche upregulate vascular support molecules such as *Vegfa* to offset TM-derived deficits. In support of this, we found an age-dependent onset of *Vegfa* expression from macrophages (starting at middle age) and supportive interactions with TM cells (Figure 2E, F), suggesting a compensatory paracrine mechanism by which immune cells may help preserve SC function in the aged outflow pathway.

### Age-dependent upregulation of immune and complement pathway genes reflects an adaptive response across TM subtypes

We investigated whether components of the innate immune response are dysregulated in the aging TM cell subtypes. Transcriptomic analysis revealed a robust and consistent age-dependent upregulation of *Lyz2* (Lysozyme 2) across all three TM subtypes (Figure 3A-D). LYZ2 is a key effector of the innate immune system and has been implicated in the modulation of inflammatory signaling, macrophage activation, and the regulation of tissue homeostasis under conditions of cellular stress^37–40^. Its broad upregulation across all TM subtypes suggests a generalized activation of innate immune programs within the TM with age, rather than a subtype-specific response. To validate this transcriptomic finding, we confirmed increased LYZ2 protein abundance in limbal strip tissue using orthogonal proteomic profiling using mass spectrometry (MS) and western blotting (Figure 3E, F). Furthermore, immunohistochemical (IHC) analysis demonstrated elevated LYZ2 immunoreactivity not only within TM cells (Figure 3G) but also across other anterior segment cell populations (sclera, for instance), suggesting that the age-associated innate immune response extends beyond the TM and may reflect a more global shift in the immune landscape of the anterior segment.

**Figure 3:**
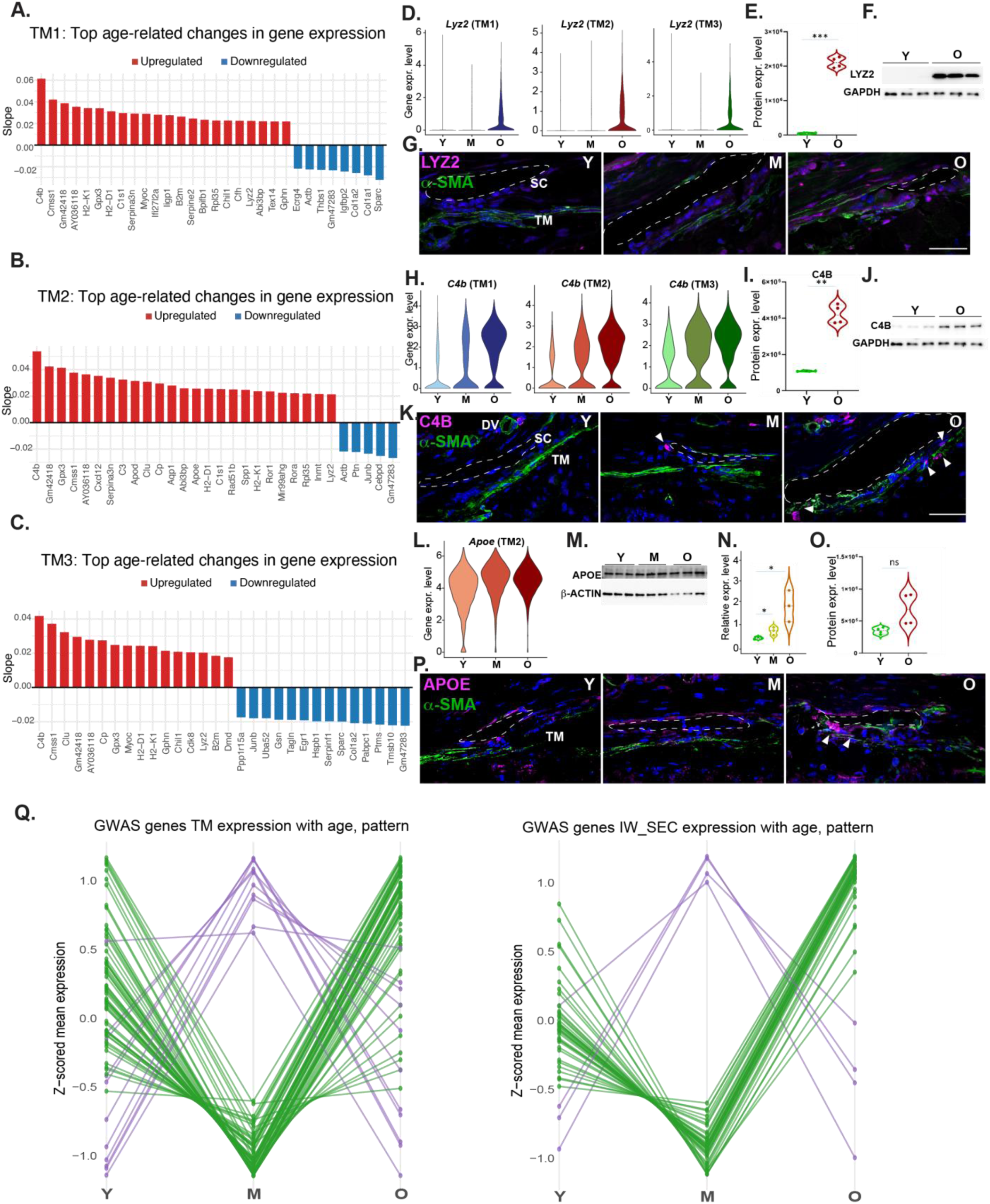
**(A-C)** Age-related gene expression changes in TM1, TM2 and TM3 cells, respectively. **(D)** Violin plot showing *Lyz2* expression in TM subtypes with age. **(E,F)** LYZ2 protein expression levels in limbal strips as detected by proteomics analysis (E, relative expression levels in 3 vs. 30-month-old mice, p < 0.0001) and Western blot analysis (F, protein levels in 3 vs. 26-month-old mice, p = 0.003). **(G)** Anterior segment sections stained with antibodies against LYZ2 and aSMA (which delineates the TM). Dashed lines mark SC. **(H)** Violin plot showing *C4b* expression in TM subtypes with age. **(I,J)** C4B protein expression levels in limbal strips as detected by proteomics analysis (**I**, relative expression levels in 3 vs. 30 month-old mice, p = 0.0013) and Western blot analysis (**J**, relative protein levels in 3 vs. 26 month-old mice, p = 0.0003). Anterior segment sections stained with antibodies against C4B and aSMA. **(K)** Anterior segment sections stained with antibodies against C4B and aSMA (which delineates the TM). Dashed lines mark SC and C4b-expressing TM cells are marked with arrows. **(L)** Violin plot showing *Apoe* expression in TM2 subtype with age. **(M-O)** APOE protein expression levels in limbal tissues as detected by Western blot analysis (**M** and **N**, relative protein levels in 3, 12 and 24 month-old mice, p = 0.04 between Y and M; p = 0.02 between Y and O) and proteomics analysis (**O**, relative expression levels in 3 vs. 30 month-old mice, p = 0.07). **(P)** Anterior segment sections stained with antibodies against APOE and aSMA (which delineates the TM). Dashed lines mark SC. **(Q)** Expression pattern of genes associated with elevated IOP and POAG in GWAS studies.

In parallel, we observed a significant age-dependent increase in the expression of *C4b* (Complement Component 4B) across all three TM subtypes (Figure 3A-C, H). C4B is a central effector molecule of the classical complement cascade. Consistent with the transcriptomic data, increased C4B protein levels were confirmed by MS and western blotting of limbal strip tissue (Figure 3 I, J). We performed limited IHC to localize C4B protein expression within the TM and confirmed that TM cells were the major site of immunoreactivity (Figure 3K), collectively providing multi-platform validation of C4B elevation in the aged TM.

We additionally identified an age-dependent upregulation of *Apoe* (Apolipoprotein E) specifically in the TM2 subtype (Figure 3L). APOE is a multifunctional lipoprotein^41,42^ and is increasingly recognized as a potent immunomodulatory molecule capable of suppressing microglial and macrophage activation, modulating complement activity, and regulating neuroinflammatory responses^43,44^, all functions of relevance in the context of the age-related immune activation observed in the TM. A recent study found that APOE was increased in the aged macaque TM and that silencing APOE in primary human TM cells in culture resulted in an increased ATP production^45^. The subtype-specific upregulation of *Apoe* in TM2 may therefore reflect a localized adaptive response aimed at modulating lipid homeostasis and tempering immune activation within this distinct TM population. Increased APOE protein abundance in TM tissue was confirmed at the protein level by western blotting and spatial localization to TM was confirmed by IHC (Figure 3M, P); MS showed a similar upward trend but did not reach statistical significance (Figure 3N, O).

### Expression trajectories of GWAS-implicated glaucoma genes suggest transcriptional priming for adaptive changes in middle age

We closely examined three age-regulated genes in TM cells - *Angptl7*, *Myoc*, and *Aqp1*, with established roles in TM biology^36,46–54^. We found that *Angptl7* (Angiopoietin-like protein 7) expression decreases with age, a finding corroborated by western blotting of ANGPTL7 in limbal strips (Figure S5A, B). However, ANGPTL7 is a secreted protein, and its expression was detected in fewer than 5% of TM cells (Figure S5A) and was also strongly expressed in corneal keratocytes (Figure S2B), precluding IHC validation on sections due to our inability to isolate TM-specific ANGPTL7 signal (Figure S5A).

While these results therefore remain inconclusive, they highlight the need for further investigation into the potential protective role of *Angptl7* in aging and glaucomatous eyes. *Myoc* (Myocilin), was found to be upregulated in TM1 and TM3 subtypes with age, alongside a similar change in protein level identified using MS and western blotting (Figure S5 D-G). *Aqp1* (Aquaporin-1), was slightly upregulated in TM cells with age at the transcript level but decreased at the protein level by western blotting (Figure S5H–J). We were unable to corroborate this using MS of limbal strips or by IHC, potentially reflecting post-transcriptional regulatory mechanisms or technical constraints.

A recent study conducted as part of the Human Cell Atlas found that many age-related genes in the TM and ciliary body begin shifting in expression between the ages of 40 and 60^55^, prior to age-related decline; a period that roughly corresponds to the 12-month time point in mice. Analysis of genes implicated in elevated IOP and POAG by GWAS revealed non-linear, age-dependent expression trajectories (Figure 3Q, S4A-D), with many genes showing significant up- or down-regulation at middle age followed by a partial reversal in expression pattern, which may reflect a transcriptional priming event, potentially representing an adaptive tissue response to counteract early aging-induced changes.

### Aging drives macrophages to the inner wall of SC

We next examined differentially expressed genes with age in IW SECs. Notably, *Selp*, a gene typically enriched in OW SECs^30,56^, showed increased expression with age in IW SECs (Figure 4A). We also noticed an increase in *Selplg* (encoding PSGL-1 that binds to P-SELECTIN) in macrophages with age (Figure 4B). Because *Selp* is commonly used as a biomarker of inflammation in aging tissues^57,58^ and mediates leukocyte tethering and persistent interactions with the endothelium, we investigated whether additional inflammatory markers were similarly upregulated in aged IW SECs. We observed increased expression of *Il6* and *Cd74*, along with MHC class I and II genes, collectively suggesting an age-related increase in immune regulation (Figure 4C). To validate these transcriptional changes at the protein level, we performed IHC for IBA1 (an established marker for macrophages^59^) and P-selectin (P-SEL) on tissue sections and found increased expression of both markers in the IW with age (Figure 4D), whereas IBA1 levels on the distal vessel side (OW, collector channels, limbal vessels) remained stable across age groups (Figure S6A, B). Western blotting of limbal strips corroborated this finding, revealing a modest but statistically significant increase in IBA1 expression with age (Figure 4E, F). Consistent with these results, GO analysis revealed an enrichment of immune regulatory pathways in aged SC relative to young, accompanied by a corresponding reduction in metabolic processes and oxidative phosphorylation pathways (Figure 4G). We examined predicted ligand-receptor interactions and found evidence of *Vegfa* signaling from macrophages to IW SECs emerging in middle-aged and persisting into old tissues that was absent in young tissues (Figure 4H). Together, these findings suggest that macrophages contribute to an adaptive compensatory response that helps maintain SC homeostasis during aging.

**Figure 4:**
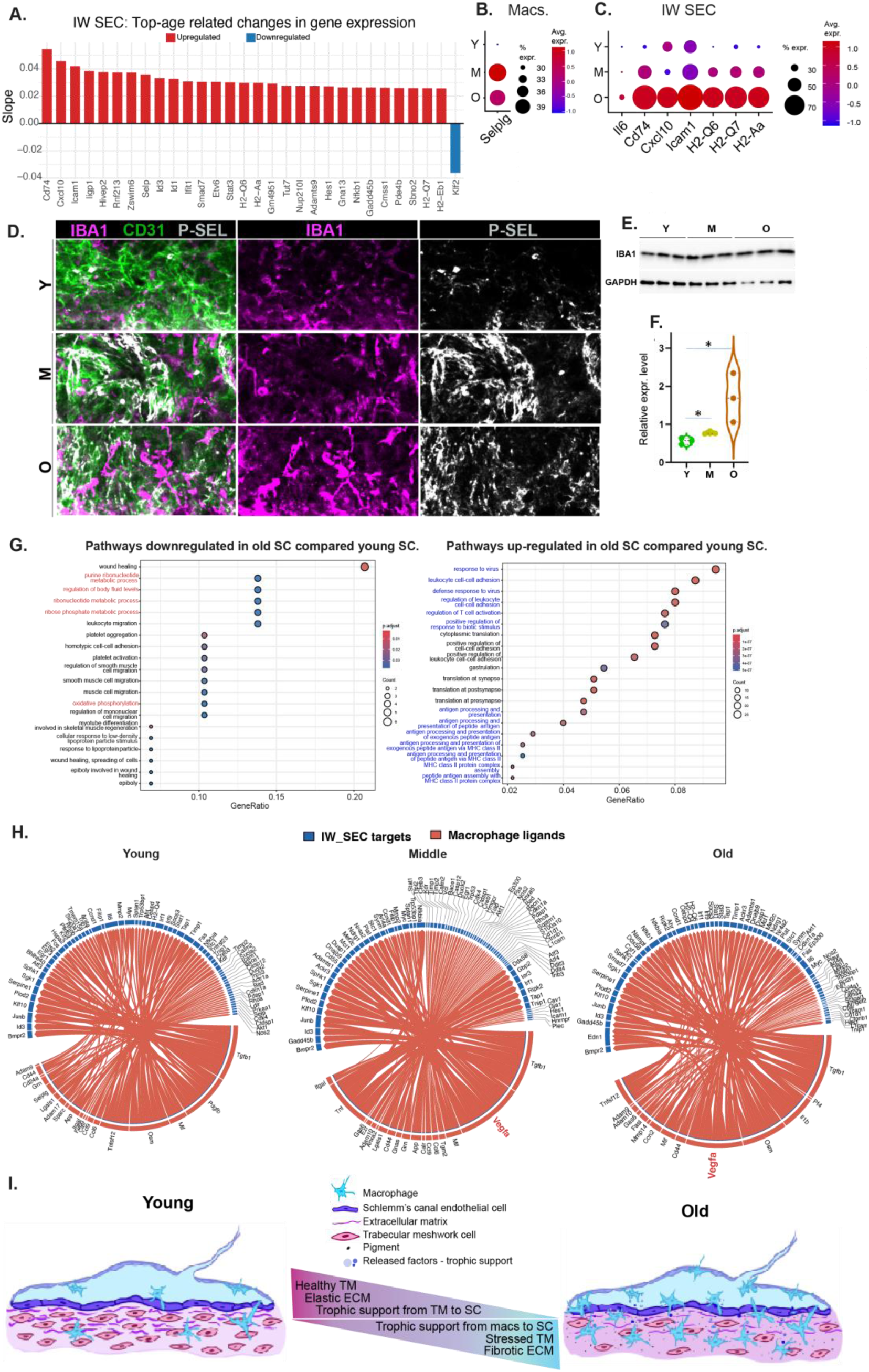
**(A)** Age related changes in gene expression of various genes within IW SECs. **(B)** Expression levels and percent cells of *Selplg* in macrophages across age groups. **(C)** Expression levels and percent cells of immune-pathway genes within IW SECs across age groups. **(D)** Anterior segment whole mounts stained for antibodies against IBA1 (macrophage marker), P-SEL (*Selp’*s corresponding protein) and CD31 (SC); **(E-F)** IBA1 protein expression levels in limbal strips as detected by Western blot analysis (relative protein levels in 3, 12 and 24 month-old mice, p = 0.01 between Y and M; p = 0.04 between Y and O). **(G)** GO analysis of pathways up and down regulated in old compared to young SC. **(H)** Circos plots showing predicted interactions between IW SEC targets and macrophage ligands across age groups. **(I)** A schematic of model for adaptive IOP homeostasis with age: Aging leads to a decrease in ECM assembly/maintenance and the number of TM cells (resulting in less trophic support from TM to SC). The amount of pigment in the TM space also increases. To compensate, macrophages infiltrate the limbic strip and release factors that provide trophic support to SC, which assist with regulation of IOP homeostasis and structural maintenance despite age-related changes and deterioration.

## Discussion

### TM/SC as a model for adaptive homeostasis with age

Despite continuous exposure to oxidative, particulate, and mechanical stress, non-glaucomatous outflow tissues function normally to maintain a stable IOP throughout life^9,60^. In the present study, we documented the changing genomic landscape of mouse outflow tissues with age, in a cell subtype-dependent fashion. TM cells show evidence of increased cellular stress, dysfunction, and increased apoptosis with age, reflected in downregulation of genes associated with ECM maintenance and trophic support for SC. Concurrent with these changes, we observed an increase in pro-inflammatory signaling from both TM and SC cells, and an increase in macrophages at the inner wall of SC and TM with age.

Significantly, as the expression of trophic mediators for SC decreased in TM cells, compensatory expression of trophic mediators like VEGFA increased in immune cells. These findings are consistent with features of inflammaging^61^. Low-grade immune activation is increasingly recognized as a hallmark of healthy aging^62^, whereby chronic innate immune activation accompanies aging across multiple tissues^63,64^. Whether these inflammatory changes are beneficial, detrimental, or context-dependent remains unclear. Macrophages are especially interesting because they are not only inflammatory cells. Tissue macrophages support surveillance, debris clearance, repair, remodeling, signaling, and local homeostasis (including in the outflow pathway)^65^. It is becoming increasingly recognized that aging tissues are not just passively deteriorating. They undergo adaptive remodeling, altered intercellular communication, chronic low-grade inflammation, and variable resilience changes^62^. Our data suggest that healthy TM/SC aging involves adaptive remodeling that preserves IOP homeostasis, with macrophages representing an important component of this resilience program. Together, our molecular and physiologic data suggest that TM/SC aging may include dynamic immune activation and remodeling to preserve physiological function.

### A midlife adaptive switch in the TM sustains outflow function into old age

With increasing age, TM cells exhibit signs of elevated cellular stress and apoptosis. Consistent with this, expression of the stress-responsive gene *Cdkn1a* (encoding p21) increased across all three TM subtypes at middle age (Figure S4A-C), followed by a subsequent decline in mRNA expression in old age. In our previous study, we reported accumulating p21 protein levels in the outflow tract (IHC) alongside reduced TM cellularity with age^14^, likely reflecting cumulative cellular responses to particle/debris accumulation, UV-induced reactive oxygen species, and mechanical stress from daily IOP fluctuations experienced by outflow cells over time. Given that p21 protein is subject to extensive post-transcriptional and post-translational regulation^66,67^, it is plausible that the middle-age induction of *Cdkn1a* transcription drives an initial rise in p21 protein that is subsequently maintained at old age. With age, a compromised TM shows evidence of a shift from active ECM protein turnover (decreased collagen expression) to fibrosis/keratosis/calcification (upregulation of keratins, *S100a4*, *Tgfβ1/Tgfβ2* and *Spp1*). These changes in the aging TM are consistent with increased tissue stiffness and contractility as measured previously^14^, and with increased risk of POAG associated with pro-fibrotic genes, particularly TGF family members by GWAS^53^. These changes promote increased stiffness of TM and SC cells, which compromises their ability to sense and respond to changes in IOP^68–70^. Consistent with this idea, results here show that *Yap1* is upregulated in old SC and TM cells (Figure S5L), possibly a compensatory change in response to increased outflow tissue stiffness.

YAP1 (Yes-associated protein 1) is a key mechanotransducer regulating TM responses to mechanical stress, and genetic variation at the YAP1 locus has been associated with POAG^36^. In cell culture, TM and SC stiffness drives YAP activation, while YAP inhibition increases outflow facility in perfused mouse eyes^71,72^.

A substantial proportion of genes associated with POAG and ocular hypertension (from GWAS studies^36,53,73–75^) unexpectedly display non-linear differential gene expression trajectories across age. Specifically, these genes show a distinct V-shaped pattern with a pronounced dip or elevation in expression at middle-age relative to young, followed by a reversal in old age. This inflection suggests a discrete priming event occurring at middle age, which we hypothesize initiates an adaptive homeostatic response, including tissue remodeling, that helps preserve TM and SC function with age.

Expression of TM genes that provide essential trophic support for SC endothelial cells was reduced coincident with age-related increases in TM stress, stiffness, and dysfunction. The largest change was observed in TM2 and included downregulation of *Vegfa* and *Svep1*. Previous studies have shown that VEGF and angiopoietin family members are essential for SC development and function^33,34,56,76–79^. In humans, gene sets associated with risk for ocular hypertension and POAG are enriched for pathways involved in lymphangiogenesis and vascular development^36^.

### Reframing inflammaging as a compensatory adaptive response in aging TM and SC

Like other aging connective tissues, the TM shows signs of senescence and expresses higher levels of inflammatory cytokines, immune modulators, growth factors, and proteases, reminiscent of a senescence-associated secretory phenotype^80^. For example, upregulation of C4b and C3 in TM subtypes implicates an age-associated activation of the complement system within the outflow pathway, which may contribute to chronic low-grade inflammation and tissue remodeling in the aging TM. Other highly upregulated proinflammatory indicators include the innate immune gene, *Lyz2*, the lipid transport gene, *Apoe,* and down-regulation of the negative regulator of monocyte activation, *Klf2*^81^. KLF2 also regulates endothelial autophagy^82^ and vasculogenesis and is itself regulated by blood flow^83^. Relevant to SC, KLF2 expression declines in various vascular diseases^84–86^. In parallel with increased expression of immune modulators by the TM, SC also significantly upregulated several immune modulatory genes with age. For example, *Il6*, *Cd74*, *Selp* and *Icam1* were among the most highly upregulated genes in the aging inner wall of SC (Figure 4A). Coincident with upregulation of these pro-inflammatory genes was an increase of immune cell abundance in outflow tissues.

Importantly, a recent study has shown that resident tissue macrophages are essential for outflow homeostasis and that resident macrophages are almost completely replaced by monocytes by 12 months of age^65^, prompting us to hypothesize that these immune cells were restoring function to an aging tissue. Consistent with this idea, we observed increased *Vegfa* expression in immune cells in outflow tissues with age, potentially compensating for age-related loss of trophic support to SC by the TM. We also see an elevated *Selp* (encodes P-selectin) in the aging IW and increase in *Selplg* in aging macrophages, which may support beneficial immune cell function. Beyond its well-established role in leukocyte rolling and adhesion, accumulating evidence indicates that sustained P-selectin/PSGL-1 signaling can also influence macrophage activation^87–89^, persistence, and tissue-remodeling programs during chronic inflammation and tissue repair. Furthermore, macrophage-derived *Mmp9* has been implicated in anchoring P-selectin at the cell surface^90^, potentially reinforcing this signaling axis. Within the context of our findings, these observations raise the possibility that sustained P-Selectin/PSGL-1 signaling contributes to establishing a persistent, tissue-supportive macrophage population that provides trophic support, including VEGFA, and promotes adaptive remodeling within the aging outflow pathway. These hypotheses now warrant direct experimental testing.

Despite a myriad of molecular and cellular changes that occur in the TM and SC with age, outflow facility and thus IOP remain stable withing a few millimeters of mercury in most people over their lifetime^9,60^. However, in 5-10% of the population (depending upon ancestry), IOP becomes dysregulated and increases with age, dramatically increasing the risk for glaucoma and blindness^6–8,91^. The present study defines the changing molecular landscape of the aging outflow pathway, revealing adaptive homeostatic mechanisms that preserve physiological outflow function while identifying vulnerabilities that may ultimately predispose to ocular hypertension and glaucoma. A companion study demonstrates that macrophage-derived VEGFA compensates for the age-related decline in TM-derived VEGFA signaling, thereby helping preserve SC homeostasis (Kiyota et al., submitted concurrently). However, the decline of additional TM-derived supportive factors, including SVEP1, suggests that macrophage-derived VEGFA alone likely cannot explain maintenance of physiological outflow function during aging, implying that additional adaptive mechanisms remain to be identified.

Collectively, these findings support a model in which aging-associated TM dysfunction promotes adaptive immune remodeling that helps preserve SC homeostasis and physiological outflow function despite lifelong insults to the outflow pathway.

## Materials and Methods

### Animals

C57BL/6J (B6) mice (males and females, ages 3–30 months) were used in this study. All mice were handled in accordance with protocols approved by the Institutional Animal Care and Use Committee of Duke University (protocol: A226-21-11-24), Columbia University (protocol: AC-AABX6652) and Ohio State University (protocol 2024A00000072) and in compliance with the Association for Research in Vision and Ophthalmology (ARVO)’s Statement for the Use of Animals in Ophthalmic and Vision Research. The mice were purchased from the Jackson Laboratory (Bar Harbor, Maine, USA), bred/housed in clear cages and kept in housing rooms at 21°C with a 12:12 h light: dark cycle.

### Limbal tissue dissections and sample preparation for single cell RNA sequencing

Columbia University protocol: Male and female mice (3, 12, and 22 months) were used. One sample was generated from a pool of 6 limbal strips (3 animals). Limbal strip was dissected as previously described^29,30^. Tissues were digested enzymatically using Papain and Deoxyribonuclease I for 20 min at 37°C and stopped using Earl’s balanced salt solution (EBSS). Cells were triturated using an 18-gauge needle, centrifuged at 300 × *g* at 4°C, washed with cold DMEM, and filtered using a 100 μm Flowmi Cell Strainer. Cells were resuspended in cold DMEM and placed on ice immediately. When the core was not available for sequencing during COVID-19 in 2020, single cells were frozen in DMEM with FBS and 10%DMSO, stored in liquid nitrogen. Cells were thawed when ready for submission, counted, and sequenced. Cells were counted using the Countess II automated cell counter. Single-cell sequencing (10 X genome) was performed at the Columbia Genome Center.

Duke University protocol: Three age groups of male mice (3, 12 and 24 months) were used. One sample/group was generated from a pool of 4-5 animals (8-10 eyes total). The eyes were dissected and strips of limbal tissue including TM/SC were collected. Single cell dissociation was performed using Collagenase IV, Dispase II and DNAse I for 60 min and then trypsin/EDTA for 10 min at 37°C (see Table S2 for details). Cells were triturated using a 1 ml pipette tip, centrifuged at 300 × g at 4°C, washed with cold DMEM containing 10% FBS, and filtered using a 70-μm cell strainer. Cells were resuspended in cold DMEM containing 5% FBS and placed on ice immediately.

### Analysis of sequencing data

Raw reads were mapped to the mm10 reference genome by 10x Genomics Cell Ranger pipeline. ‘Seurat’^92^ (version 4.0.1) was used to conduct all single-cell and single-nucleus sequencing analyses. Briefly, the dataset was filtered to contain cells with at least 200 expressed genes and genes with expression in more than three cells. Cells were also filtered for mitochondrial gene expression (<20%). The dataset was log-normalized and scaled. Biological incompatibility based on gene expression was used to identify doublets. Soupx was used to remove ambient RNA contamination^93^. For the scRNA-seq dataset generated at Columbia, we performed unsupervised clustering and compared the results to previous annotations^30–32,34,94^. Cluster identities were assigned based on a combination of known marker genes and validation experiments. For the Duke scRNA-seq dataset, unsupervised clustering identified clusters like those found in the Columbia dataset. When we integrated the two datasets, the integration was cogent and did not produce a separate, dataset-specific cluster, indicating congruence in gene expression across the two datasets. All subsequent analyses were performed on this integrated dataset.

#### GO analysis

Gene ontology and pathway enrichment analysis were performed by comparing the differentially expressed genes in young versus old samples. For comparisons of all TM cells to all sequenced cells, pathway enrichment was compared against a background expression universe of genes expressed in any sequenced cell. For comparisons of individual TM cell subtypes, only genes expressed in TM cells were used as a background (p value cut-off 0.01)^95^. Pathways that were significantly enriched or underrepresented were also analyzed (P value cut-off 0.01). The R package ClusterProfiler was used for analysis^96^.

#### Predicted ligand-receptor interactions

Predicted ligand-target links between interacting cells were identified using LRLoop^97^, which was developed based on NicheNet^98^. Briefly, the expression of genes in cell types is linked to a database of signaling and gene regulatory networks curated from prior information, enabling viable predictions of potential interactions between cell types. These interactions were calculated using target molecules from one cell type and ligands from multiple cell types. The top interactions are represented in a Circos plot.

#### Age-associated expression in TM and SEC subcluster

Age-associated transcriptional changes were assessed within the TM and (inner wall Schlemm’s endothelial cells) IW SEC subclusters independently. Gene expression was first modeled as a function of age using linear regression. Genes expressed in at least 25% of aged cells were then retained, and genes demonstrating significant (absolute slope > 0.001, P-value < 0.05) age-associated expression changes were ranked by the magnitude of their regression coefficient. The top thirty genes with the largest absolute age-associated slopes were visualized using waterfall plots, with each bar representing the regression slope of an individual gene.

#### Age-dependent trajectory of GWAS genes in TM and IW SEC

To evaluate age-dependent expression patterns of glaucoma-associated genes, GWAS-identified genes were analyzed within the TM cluster and IW SEC subcluster. Mean expression values were calculated for each gene across young, middle-aged, and old groups and standardized using z-score normalization to compare relative expression trajectories. Expression trajectories were visualized using line plots, showing dynamic changes with aging.

### Proteomics

Two age groups of mice (3 and 30 months old) were used and limbal strips from 4 eyes/2 mice were pooled for each sample (N=2, total 4 mice/8 eyes for each age group). Limbal strips were carefully dissected on ice and the tissues were collected and frozen at −80°C. Proteins were isolated using lysis buffer 2% SDS, 0.1M Tris-HCl pH8 and proteinase inhibitor. Protein lysates (20 μg/sample) were solubilized, reduced, alkylated and subjected to tryptic hydrolysis using SP3 protocol (PMID: 25358341). The protein digests were separated by liquid chromatography using the Q Exactive HF Orbitrap mass spectrometer (Thermo Fisher Scientific). For protein identification and label-free relative protein quantification, raw mass spectral data files were imported into Progenesis QI for Proteomics 4.2 software (Nonlinear Dynamics) for duplicate run alignment of each preparation and peak area calculations. Only proteins identified on the basis of 2 or more peptides (protein confidence P < 0.05 and false discovery rate < 1%) were included in the protein quantification analysis.

### Western blot (WB) analysis

Protein lysates were prepared in the same manner as for proteomics and separated via SDS-PAGE and transferred to nitrocellulose membranes. Blots were cut into three parts: up, middle and low. C4b (∼190kDa), Lyz2 (∼14kDa), AQP1 (∼28kDa), Angptl7 (∼45kDa), Myoc (∼55-57kDa) and GAPDH (∼37kDa). Limbal tissues from one mouse/two eyes were used per sample, experiments were repeated three times for each age group). Membranes were blocked and probed with respective primary and secondary antibodies. Immunoblots were developed using SuperSignal West Femto Maximum sensitivity substrate (Thermo Fisher Scientific), followed by scanning and analysis using ChemiDoc Touch imaging and Image Lab Touch Software (Bio-Rad Laboratories). The intensities of specific protein bands were normalized by GAPDH or beta actin and quantitatively analyzed using ImageJ software.

### Statistical analysis

Western blot and proteomics data were analyzed using one-way ANOVA and/or Welch’s *t* test. P < 0.05 indicated significant difference.

### Immunohistochemistry (IHC)

Eyes from 3, 12 and 22-24 months old mice were collected and fixed in 4% paraformaldehyde at 4°C overnight.

#### For sections

At Duke University, eyes were bisected, and the posterior segments and lenses were removed. The anterior segments were cut into four quadrants. One quadrant was embedded in optimal cutting temperature (OCT) embedding media (Tissue-Tek) and cut into 10-μm-thick sections using a CM 1950 Cryostat (Leica Biosystems). At Columbia University, eyes were processed by making a window to the back of the eye cup by removing the optic nerve. Eyes were equilibrated in 30% sucrose and embedded in OCT. Eyes were stored at –80 °C until they were sectioned and processed for immunostaining. Sections were briefly washed in PBS and 0.3% PBST and blocked with blocking buffer made of 10% donkey serum in PBST. Primary antibodies were applied in blocking buffer overnight at 4 °C and secondary antibodies with DAPI for 2 hr at room temperature. Slides were washed and cover-slipped. All sections were processed at the same time. Staining was performed on three eyes per age group, and 4-5 sections/slide were analyzed.

#### For whole mounts

The anterior segment including the limbus was dissected in ice-cold PBS. The iris was carefully removed and shallow cuts were made to the anterior eye cup to display four quadrants in a petal-like pattern. The tissue was incubated in blocking solution of 0.3% PBST with 3% BSA for 2 hr at room temperature. Primary antibodies in blocking solution were added and the eyes left at 4 °C for 2–3 days. After four PBST washes of an hour each at room temperature, secondary antibodies along with DAPI were added and eyes placed in the dark at 4 °C overnight. After four PBS washes of an hour each at room temperature, eyes were mounted on a slide, cover-slipped, and imaged. Staining was performed on three eyes per age group, and 8 images per eye were analyzed.

Immunostaining was performed with specific antibodies against Iba1, C3, C4b, Apoe, YAP1, Lyz2, KLF2, aSMA, Angptl7, Myoc, Cd31, Selp, GAPDH and the corresponding secondary antibodies (see Table S1).

### Imaging and image analysis

At Columbia University, images were captured using a Leica SP8 confocal. Section images were collected at identical laser powers, gain and exposure for each antibody staining across ages. Whole mount images were captured using Leica SP8 confocal with a Z-stack of 1uM thickness (going from outer vascular bed to inner TM with SC in between). Images were rendered and processed in IMARIS. At Ohio State University, anterior segment whole mounts stained with antibodies endomucin and IBA1. Confocal Z stacks of the limbal region including the AQH draiange strucutres were captured at 0.32µm intervals using a Zeiss 900 LSM Airy Scan 2 with a 40X NA 1.1 40x LD C-Apochromat objective. The Z stacks were rendered in 3D using IMARIS and a surface was rendered using the SC channel. IBA1 positive macrophage within 2-3µm above (limbal vessel side) and below (TM side) of the SC surfaces were counted using aster rendering the IBA1 channel using Spots function in IMARIS. At Duke University, images were captured using a Nikon Eclipse 90i confocal laser scanning microscope. Images were collected at identical intensity and gain settings on the same day for each antibody staining for all three age groups.

## Funding acknowledgements

This project was supported by the BrightFocus Foundation grant CG2020004 (to SWMJ, DS and KK), BrightFocus Foundation National Glaucoma Research grant G2021007S (RB), HHMI, the Precision Medicine Initiative at Columbia University, National Eye Institute (NEI) grant R01EY032507, and the New York Fund for Innovation in Research and Scientific Talent NYFIRST – EMPIRE CU19-2660 (SWMJ). Partial support was provided by NEI grants R01EY018606 (SWMJ), R01EY032062 (SWMJ, and KK), R01EY011721 (SWMJ), R01EY030124 (DS), R01EY028608 (DS), R01EY022359 (DS), and The Glaucoma Foundation (SWMJ). Support also came from start-up funds from Columbia University (RB, SWMJ), and Ohio State University (KK). Additional support was provided by Research to Prevent Blindness (RPB) Career Development Award (RB), The Glaucoma Foundation’s grant-in-aid (RB), Glaucoma Research Foundation Shaffer research grant (RB),, and the Knights Templar Eye Foundation career starter grant (AH), RPB new chair challenge grant to Ohio State University, and unrestricted departmental award from RPB to Columbia University and Duke university (RB, SWMJ, DS). Core grant support also contributed: P30EY019007 (Columbia University), P30EY005722 (Duke university). RB is a Chang Burch Scholar. SWMJ is the Robert Burch III Professor of Ophthalmic Sciences and was an Investigator with the Howard Hughes Medical Institute (HHMI) during the first years of this project. The content is solely the responsibility of the authors and does not necessarily represent the official views of the National Institutes of Health.

---

**Table S1.** List of antibodies used in the study.

| **Antibody** | **Dilution** | **Source** | **ID** |
| --- | --- | --- | --- |
| aSMA | Sections (1:200) | Abcam | ab5694 |
| MYOC | Sections (1:200)<br>WB (1:1000) | R&D systems | AF2537 |
| PECAM (CD31) | WM (1:50)<br>Sections (1:200) | BD Pharmingen | 550274 |
| SELP | WM (1:50)<br>Sections (1:100) | R&D Systems | AF737-SP |
| VECAD | WM (1:50)<br>Sections (1:200) | R&D Systems | AF1002 |
| IBA | Sections (1:400)<br>WB (1:1000) | Wako | 019-19741 |
| C3 | Sections (1:200)<br>WB (1:500) | Invitrogen | PA1-29715 |
| C4B | Sections (1:50)<br>WB (1:1000) | Invitrogen | PA5150199 |
| APOE | Sections (1:100)<br>WB (1:200) | Invitrogen | 701241 |
| YAP1 | Sections (1:50)<br>WB (1:1000) | Cell Signaling | 14074s |
| LYZ2 | Sections (1:100)<br>WB (1:1000) | NSJ Bioreagents | FY12064 |
| ANGPTL7 | Sections (1:50)<br>WB (1:3000) | Proetientech | 10396-1-ap |
| AQP1 | WB (1:1000) | Homemade |  |

**Table S2.** List of other resources and reagents.

| Resource | Source | ID |
| --- | --- | --- |
| **Animals** |  |  |
| C57BL/6J mice | Jackson Labs | IMSR_JAX:000664 |
| **Reagents/Chemicals/Other** |  |  |
| 4% PFA | Thermo Fisher Scientific | Cat no. 50-980-495 |
| Bovine Serum Albumin | Thermo Fisher Scientific | Cat no. AM2616 |
| Collagenase Type 4 | Worthington Biochemical | Cat no. LS004188 |
| **Dispase II** | Sigma | Cat no. D4693 |
| DAPI | Thermo Fisher Scientific | Cat no. 62248 |
| Dulbecco's Modified Eagle Medium (DMEM) | Thermo Fisher Scientific | Cat no. 11965-118 |
| Dulbecco's Phosphate Buffered Saline (PBS) | Sigma Aldrich | Cat no. D8662 |
| Earle's balanced salt solution (EBSS) | Thermo Fisher Scientific | Cat no. 24010-043 |
| Papain Dissociation System | Worthington Biochemical | Cat no. LK003153 |

**S1:**
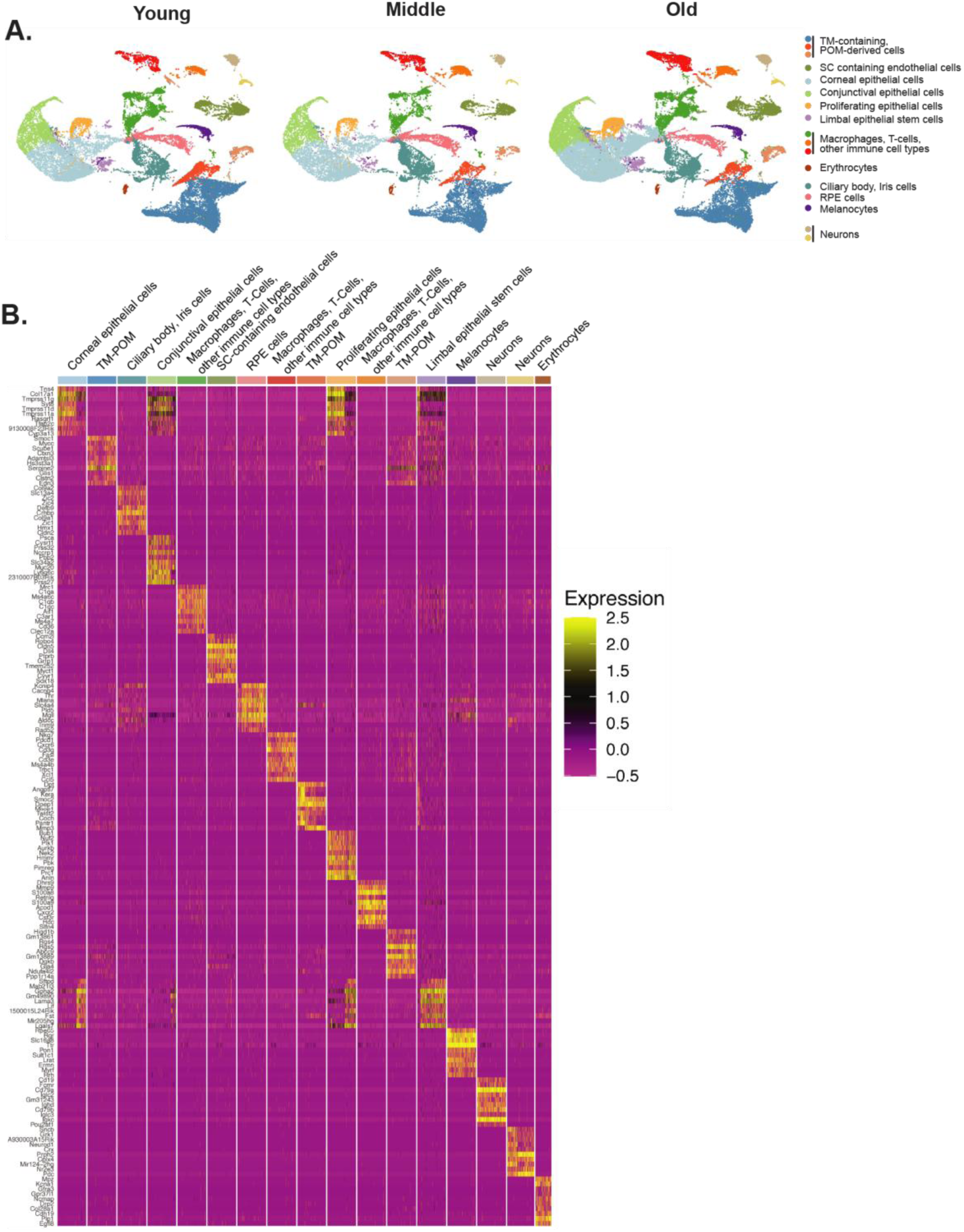
**(A)** UMAP of all major anterior segment cell types, separated by age groups. **(B)** Heat map showing expression of various genes by various cell types identified by UMAP clustering.

**S2:**
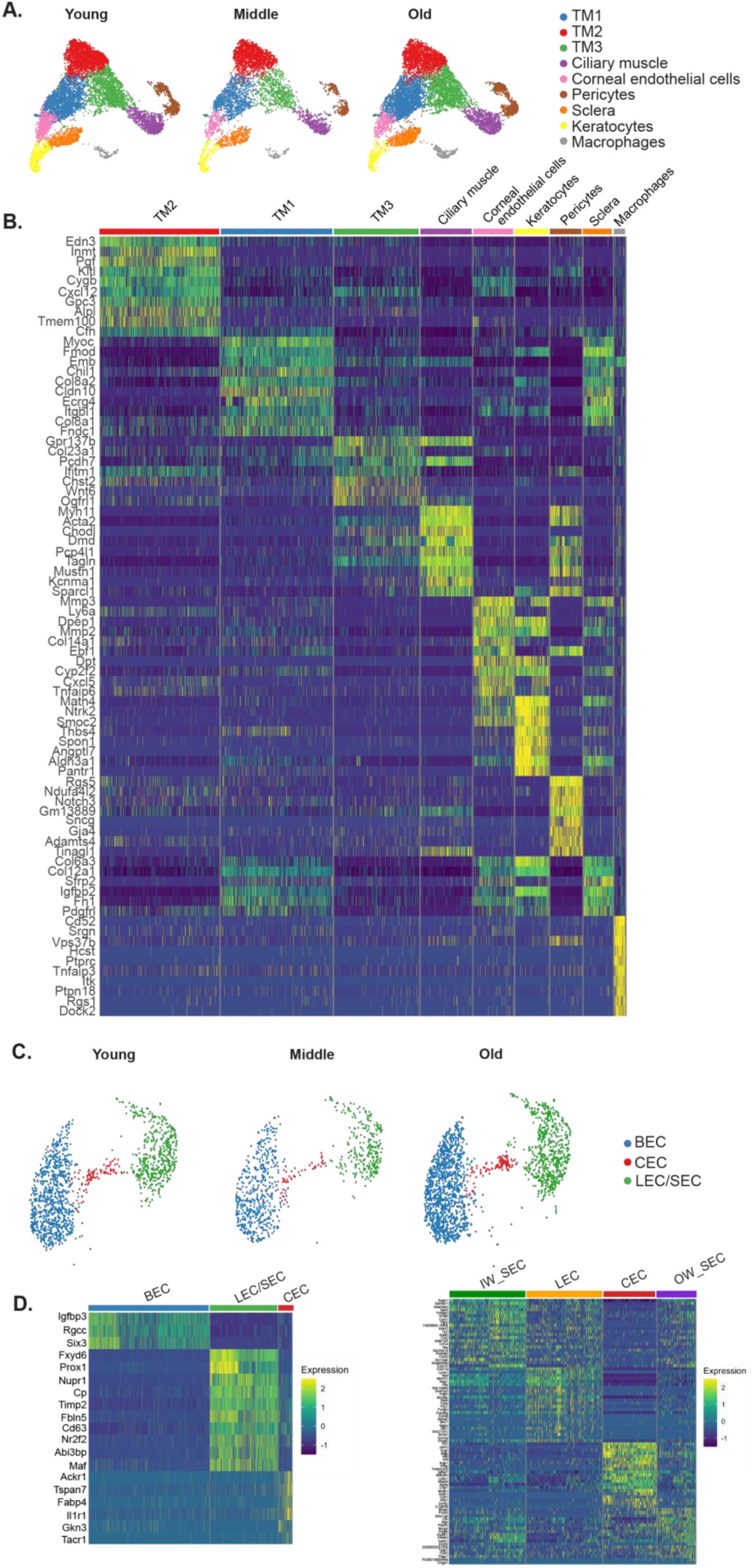
**(A)** Iterative subclustering of TM-containing POM-derived cell clusters within each age group. **(B)** Heat map showing signature genes from cell types identified by subclustering of TM-containing POM-derived cell clusters. **(C)** Iterative subclustering of SC-containing endothelial cells populations separated by age groups. **(D)** Heat map showing signature genes for BEC, LEC/SEC and CEC cells. **(E)** Heat map showing signature genes for IW SECs, LECs, CECs and OW SECs.

**S3:**
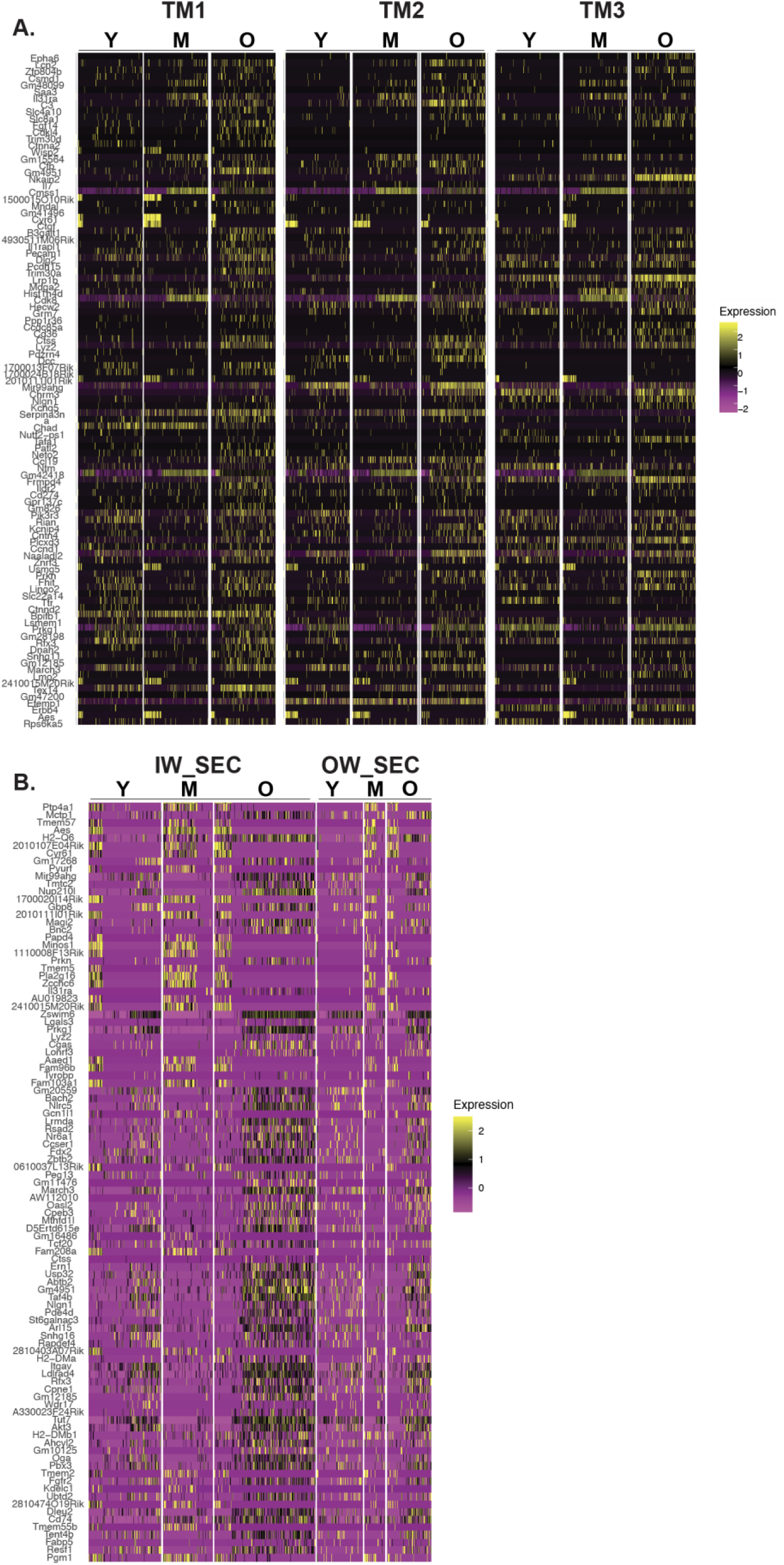
**(A)** GO analysis of gene expression in TM subtypes across ages. **(B)** GO analysis of gene expression in IW and OW SECs across ages.

**S4:**
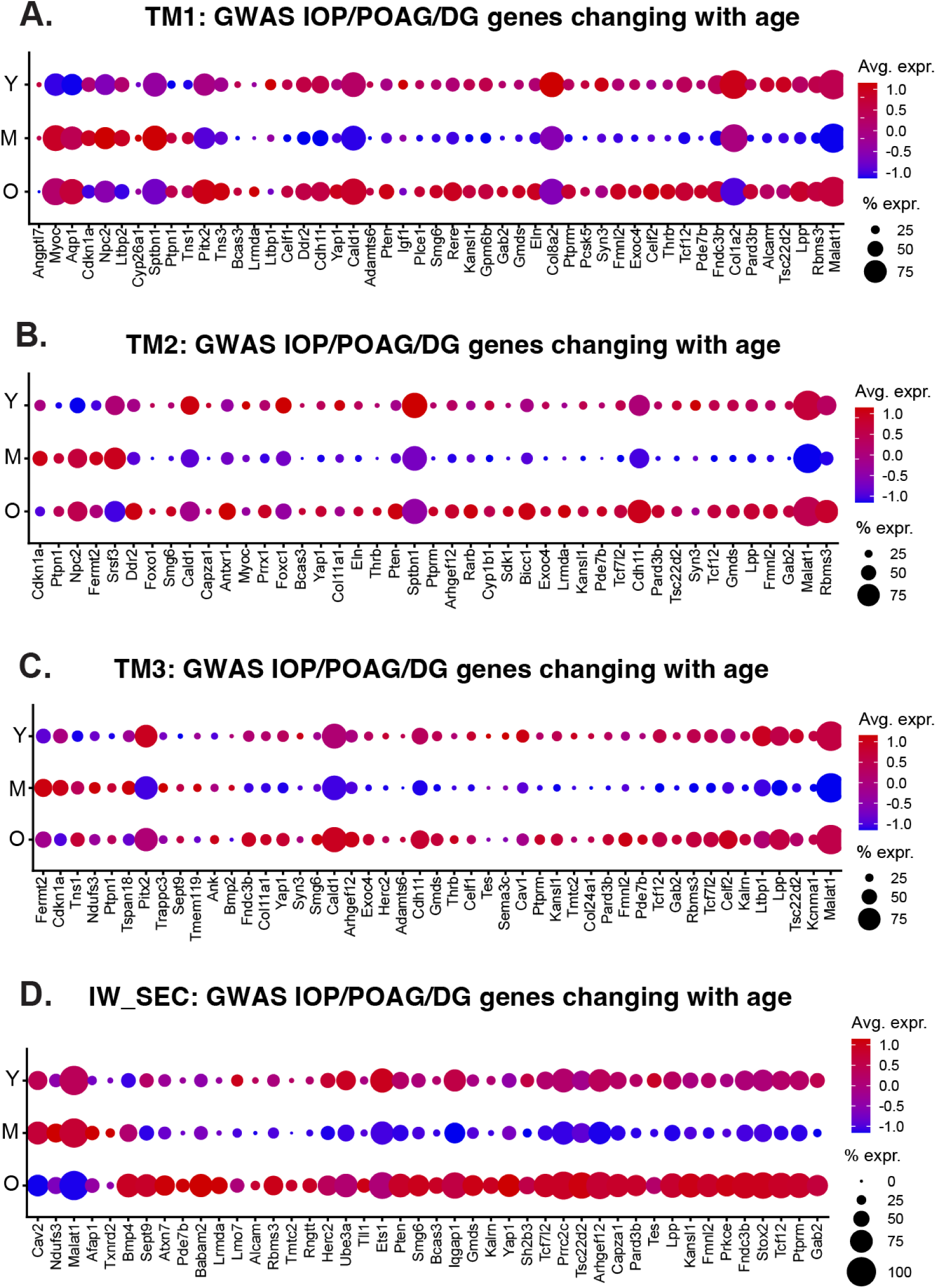
**(A-D)** GWAS analysis of age-related changes in genes associated with IOP elevation and POAG in the 3 TM cell subtypes and IW SECs.

**S5:**
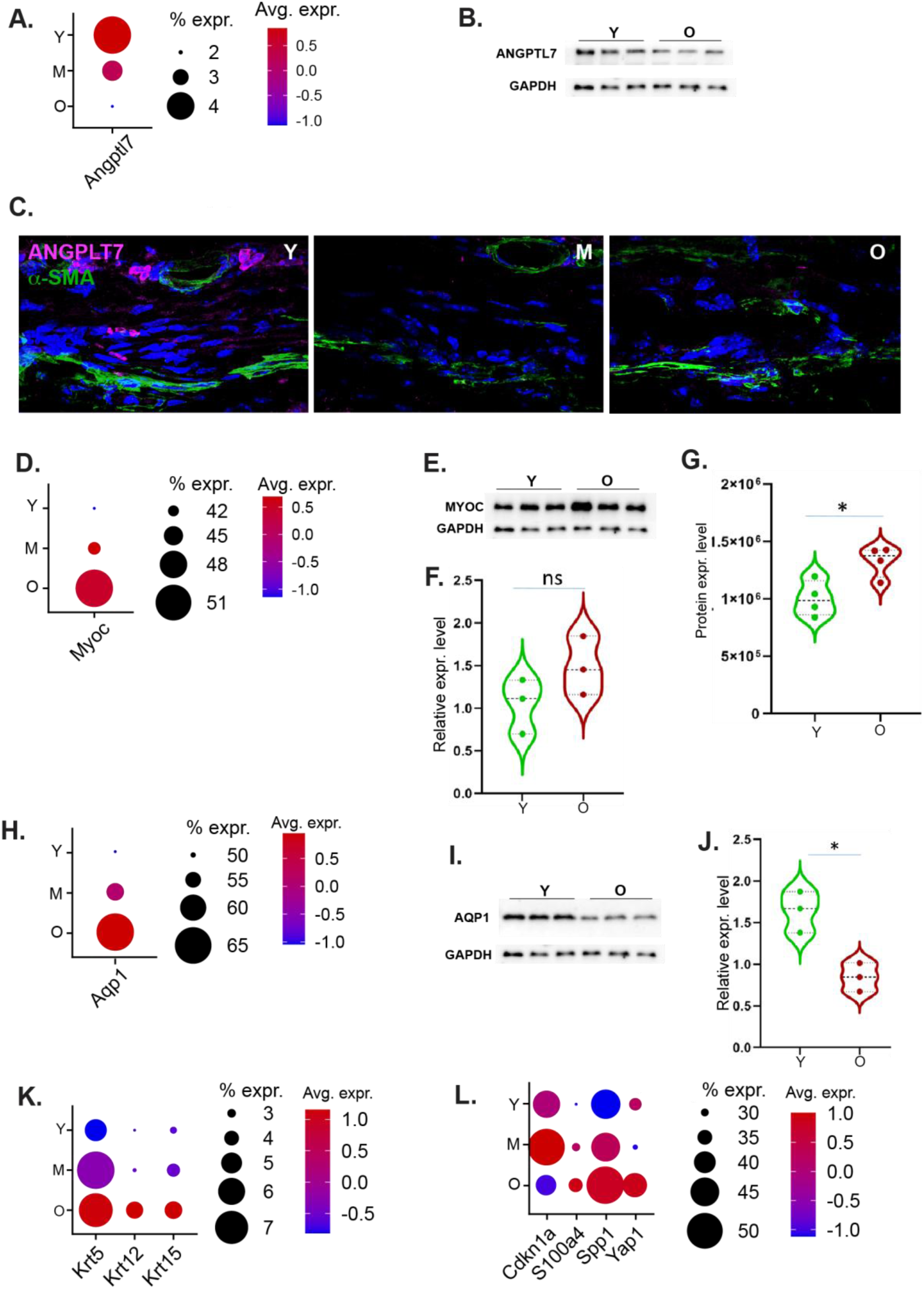
**(A)** *Angptl7* expression in TM cells with age. **(B)** ANGPTL7 protein expression levels in limbal strips as detected by Western blot analysis (3 vs. 26 month-old mice, p = 0.04). **(C)** Anterior segment sections stained with antibodies against ANGPTL7 and aSMA (which delineates the TM). **(D)** *Myoc* expression in TM cells with age. **(E-G)** MYOC protein expression levels in limbal strips as detected by Western blot analysis (**E** and **F**, protein levels in 3 vs. 26 month-old mice, p = 0.18) and proteomics analysis (**G**, relative expression levels in 3 vs. 30 month-old mice, p = 0.02). **(H)** *Aqp1* expression in TM cells with age. **(I-J)** AQP1 protein expression levels in limbal strips as detected by Western blot analysis (3 vs. 26 month-old mice, p = 0.01). **(K)** Keratin genes’ expression in TM cells with age. **(L)** Cell stress and other genes’ expression in SC cells with age.

**S6:**
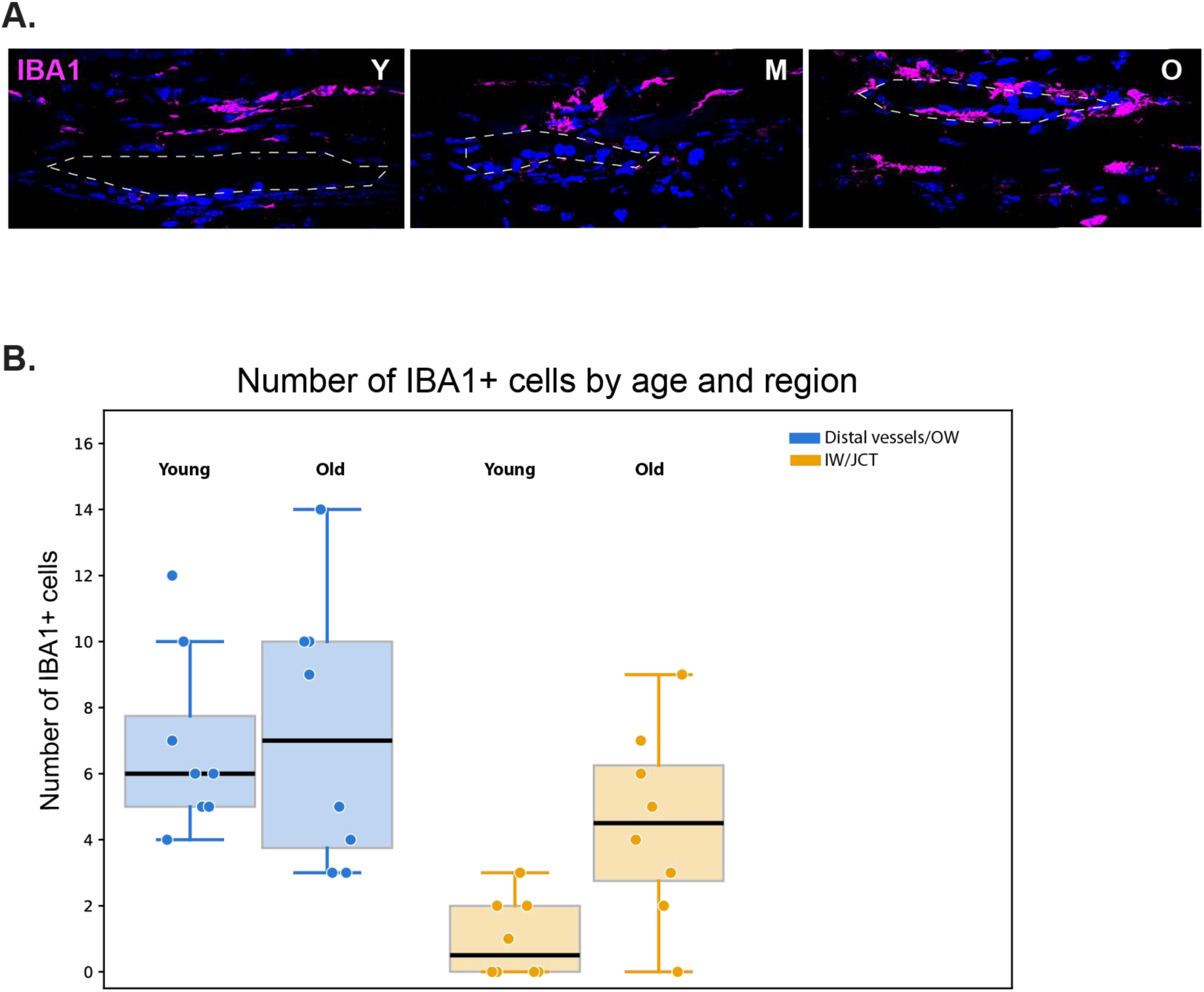
**(A)** Anterior segment whole mounts probed via IHC for IBA1 (macrophage marker). Dashed line marks SC. **(B)** Barplot comparing the number of IW/OW IBA1+ cells in the young vs old group.

## REFERENCES

1. McMonnies CW. Glaucoma history and risk factors. J Optom. 2017;10(2):71–78. doi:10.1016/j.optom.2016.02.003

2. Wiggs JL, Pasquale LR. Genetics of glaucoma. Hum Mol Genet. 2017;26(R1):R21–R27. doi:10.1093/hmg/ddx184

3. Križaj D. What is glaucoma? In: Kolb H, Fernandez E, Jones B, Nelson R, eds. Webvision: The Organization of the Retina and Visual System. University of Utah Health Sciences Center; 1995. Accessed July 6, 2026. http://www.ncbi.nlm.nih.gov/books/NBK543075/

4. Lee SSY, Mackey DA. Glaucoma - risk factors and current challenges in the diagnosis of a leading cause of visual impairment. Maturitas. 2022;163:15–22. doi:10.1016/j.maturitas.2022.05.002

5. Acott TS, Kelley MJ, Keller KE, et al. Intraocular pressure homeostasis: maintaining balance in a high-pressure environment. J Ocul Pharmacol Ther. 2014;30(2-3):94–101. doi:10.1089/jop.2013.0185

6. Gabelt BT, Kaufman PL. Changes in aqueous humor dynamics with age and glaucoma. Prog Retin Eye Res. 2005;24(5):612–637. doi:10.1016/j.preteyeres.2004.10.003

7. Toris CB, Yablonski ME, Wang YL, Camras CB. Aqueous humor dynamics in the aging human eye. Am J Ophthalmol. 1999;127(4):407–412. doi:10.1016/s0002-9394(98)00436-x

8. David R, Zangwill L, Stone D, Yassur Y. Epidemiology of intraocular pressure in a population screened for glaucoma. Br J Ophthalmol. 1987;71(10):766–771. doi:10.1136/bjo.71.10.766

9. Klein BE, Klein R, Linton KL. Intraocular pressure in an American community. The Beaver Dam Eye Study. Invest Ophthalmol Vis Sci. 1992;33(7):2224–2228.

10. Keller KE, Peters DM. Pathogenesis of glaucoma: Extracellular matrix dysfunction in the trabecular meshwork-A review. Clin Exp Ophthalmol. 2022;50(2):163–182. doi:10.1111/ceo.14027

11. Kim J, Kang JH, Wiggs JL, et al. Does Age Modify the Relation Between Genetic Predisposition to Glaucoma and Various Glaucoma Traits in the UK Biobank? Invest Ophthalmol Vis Sci. 2025;66(2):57. doi:10.1167/iovs.66.2.57

12. Stamer WD, Acott TS. Current understanding of conventional outflow dysfunction in glaucoma. Curr Opin Ophthalmol. 2012;23(2):135–143. doi:10.1097/ICU.0b013e32834ff23e

13. Carreon T, van der Merwe E, Fellman RL, Johnstone M, Bhattacharya SK. Aqueous outflow - A continuum from trabecular meshwork to episcleral veins. Progress in Retinal and Eye Research. 2017;57:108–133. doi:10.1016/j.preteyeres.2016.12.004

14. Li G, van Batenburg-Sherwood J, Safa BN, et al. Aging and intraocular pressure homeostasis in mice. Aging Cell. 2024;23(7):e14160. doi:10.1111/acel.14160

15. Lei Y, Overby DR, Boussommier-Calleja A, Stamer WD, Ethier CR. Outflow physiology of the mouse eye: pressure dependence and washout. Invest Ophthalmol Vis Sci. 2011;52(3):1865–1871. doi:10.1167/iovs.10-6019

16. Overby DR, Bertrand J, Schicht M, Paulsen F, Stamer WD, Lütjen-Drecoll E. The Structure of the Trabecular Meshwork, Its Connections to the Ciliary Muscle, and the Effect of Pilocarpine on Outflow Facility in Mice. Invest Ophthalmol Vis Sci. 2014;55(6):3727–3736. doi:10.1167/iovs.13-13699

17. Liu B, McNally S, Kilpatrick JI, Jarvis SP, O’Brien CJ. Aging and ocular tissue stiffness in glaucoma. Surv Ophthalmol. 2018;63(1):56–74. doi:10.1016/j.survophthal.2017.06.007

18. Morgan JT, Raghunathan VK, Chang YR, Murphy CJ, Russell P. The intrinsic stiffness of human trabecular meshwork cells increases with senescence. Oncotarget. 2015;6(17):15362–15374. doi:10.18632/oncotarget.3798

19. Quigley HA. Glaucoma. Lancet. 2011;377(9774):1367–1377. doi:10.1016/S0140-6736(10)61423-7

20. McDowell CM, Kizhatil K, Elliott MH, et al. Consensus Recommendation for Mouse Models of Ocular Hypertension to Study Aqueous Humor Outflow and Its Mechanisms. Invest Ophthalmol Vis Sci. 2022;63(2):12. doi:10.1167/iovs.63.2.12

21. Aihara M, Lindsey JD, Weinreb RN. Experimental mouse ocular hypertension: establishment of the model. Invest Ophthalmol Vis Sci. 2003;44(10):4314–4320. doi:10.1167/iovs.03-0137

22. Boussommier-Calleja A, Bertrand J, Woodward DF, Ethier CR, Stamer WD, Overby DR. Pharmacologic manipulation of conventional outflow facility in ex vivo mouse eyes. Invest Ophthalmol Vis Sci. 2012;53(9):5838–5845. doi:10.1167/iovs.12-9923

23. Li G, Farsiu S, Chiu SJ, et al. Pilocarpine-induced dilation of Schlemm’s canal and prevention of lumen collapse at elevated intraocular pressures in living mice visualized by OCT. Invest Ophthalmol Vis Sci. 2014;55(6):3737–3746. doi:10.1167/iovs.13-13700

24. Millar JC, Clark AF, Pang IH. Assessment of aqueous humor dynamics in the mouse by a novel method of constant-flow infusion. Invest Ophthalmol Vis Sci. 2011;52(2):685–694. doi:10.1167/iovs.10-6069

25. Overby DR, Bertrand J, Tektas OY, et al. Ultrastructural changes associated with dexamethasone-induced ocular hypertension in mice. Invest Ophthalmol Vis Sci. 2014;55(8):4922–4933. doi:10.1167/iovs.14-14429

26. Li G, Lee C, Agrahari V, et al. In vivo measurement of trabecular meshwork stiffness in a corticosteroid-induced ocular hypertensive mouse model. Proceedings of the National Academy of Sciences. 2019;116(5):1714–1722. doi:10.1073/pnas.1814889116

27. Coulon SJ, Schuman JS, Du Y, Bahrani Fard MR, Ethier CR, Stamer WD. A novel glaucoma approach: Stem cell regeneration of the trabecular meshwork. Prog Retin Eye Res. 2022;90:101063. doi:10.1016/j.preteyeres.2022.101063

28. Stamer WD, Clark AF. The many faces of the trabecular meshwork cell. Experimental Eye Research. 2017;158:112–123. doi:10.1016/j.exer.2016.07.009

29. Tolman N, Li T, Balasubramanian R, et al. Single-cell profiling of trabecular meshwork identifies mitochondrial dysfunction in a glaucoma model that is protected by vitamin B3 treatment. Elife. 2026;14:RP107161. doi:10.7554/eLife.107161

30. Balasubramanian R, Kizhatil K, Li T, et al. Transcriptomic profiling of Schlemm’s canal cells reveals a lymphatic-biased identity and three major cell states. bioRxiv: The Preprint Server for Biology. Preprint posted online August 6, 2024:2023.08.31.555823. doi:10.1101/2023.08.31.555823

31. van Zyl T, Yan W, McAdams A, et al. Cell atlas of aqueous humor outflow pathways in eyes of humans and four model species provides insight into glaucoma pathogenesis. Proc Natl Acad Sci U S A. 2020;117(19):10339–10349. doi:10.1073/pnas.2001250117

32. Patel G, Fury W, Yang H, et al. Molecular taxonomy of human ocular outflow tissues defined by single-cell transcriptomics. Proc Natl Acad Sci U S A. 2020;117(23):12856–12867. doi:10.1073/pnas.2001896117

33. Balasubramanian R, Tolman N, Li T, et al. Single-cell characterization of anterior segment development in the mouse reveals the cell types, pathways, and signals driving formation of the trabecular meshwork and Schlemm’s canal. Elife. 2026;15:RP109230. doi:10.7554/eLife.109230

34. Thomson BR, Liu P, Onay T, et al. Cellular crosstalk regulates the aqueous humor outflow pathway and provides new targets for glaucoma therapies. Nat Commun. 2021;12:6072. doi:10.1038/s41467-021-26346-0

35. Young TL, Whisenhunt KN, Jin J, et al. SVEP1 as a Genetic Modifier of TEK-Related Primary Congenital Glaucoma. Invest Ophthalmol Vis Sci. 2020;61(12):6. doi:10.1167/iovs.61.12.6

36. Gharahkhani P, Jorgenson E, Hysi P, et al. Genome-wide meta-analysis identifies 127 open-angle glaucoma loci with consistent effect across ancestries. Nat Commun. 2021;12(1):1258. doi:10.1038/s41467-020-20851-4

37. Yan HZ, Fang YW, Zhou SY, et al. Dual role of Lyz2-positive myeloid cells in traumatic brain injury: acute anti-inflammatory effects vs. chronic neurological deterioration. Front Cell Neurosci. 2025;19:1642410. doi:10.3389/fncel.2025.1642410

38. Latorre J, Lluch A, Ortega FJ, et al. Adipose tissue knockdown of lysozyme reduces local inflammation and improves adipogenesis in high-fat diet-fed mice. Pharmacol Res. 2021;166:105486. doi:10.1016/j.phrs.2021.105486

39. Kitchen-Goosen SM, Schumacher H, Good J, et al. Endometrial hyperplasia with loss of APC in a novel population of Lyz2-expressing mouse endometrial epithelial cells. Carcinogenesis. 2023;44(1):54–64. doi:10.1093/carcin/bgac101

40. Fan C, Song S, Han Y, et al. Targeting lysozyme 2 in endocardium promotes rapid recovery by modulating remote injury signals. Cell Stem Cell. 2025;32(10):1563–1576.e11. doi:10.1016/j.stem.2025.08.015

41. Liu C, Liu J, Wang YY, Xu SF, Yu LM. APOE Lipoprotein Particles: Pathophysiology, Therapy, and the Crosstalk in Alzheimer’s Disease and Cardiovascular Disease. Mol Neurobiol. 2026;63(1):325. doi:10.1007/s12035-025-05629-3

42. Yang LG, March ZM, Stephenson RA, Narayan PS. Apolipoprotein E in lipid metabolism and neurodegenerative disease. Trends Endocrinol Metab. 2023;34(8):430–445. doi:10.1016/j.tem.2023.05.002

43. Lee S, Devanney NA, Golden LR, et al. APOE modulates microglial immunometabolism in response to age, amyloid pathology, and inflammatory challenge. Cell Rep. 2023;42(3):112196. doi:10.1016/j.celrep.2023.112196

44. Yin C, Ackermann S, Ma Z, et al. ApoE attenuates unresolvable inflammation by complex formation with activated C1q. Nat Med. 2019;25(3):496–506. doi:10.1038/s41591-018-0336-8

45. Wu J, Wang C, Sun S, et al. Single-cell transcriptomic Atlas of aging macaque ocular outflow tissues. Protein Cell. 2024;15(8):594–611. doi:10.1093/procel/pwad067

46. Brown SFK, Nguyen H, Mzyk P, et al. ANGPTL7 and Its Role in IOP and Glaucoma. Invest Ophthalmol Vis Sci. 2024;65(3):22. doi:10.1167/iovs.65.3.22

47. Tanigawa Y, Wainberg M, Karjalainen J, et al. Rare protein-altering variants in ANGPTL7 lower intraocular pressure and protect against glaucoma. PLoS Genet. 2020;16(5):e1008682. doi:10.1371/journal.pgen.1008682

48. Aboobakar IF, Collantes ERA, Hauser MA, Stamer WD, Wiggs JL. Rare protective variants and glaucoma-relevant cell stressors modulate Angiopoietin-like 7 expression. Hum Mol Genet. 2023;32(15):2523–2531. doi:10.1093/hmg/ddad083

49. Resch ZT, Fautsch MP. Glaucoma-associated myocilin: a better understanding but much more to learn. Exp Eye Res. 2009;88(4):704–712. doi:10.1016/j.exer.2008.08.011

50. Tamm ER. Myocilin and glaucoma: facts and ideas. Prog Retin Eye Res. 2002;21(4):395–428. doi:10.1016/s1350-9462(02)00010-1

51. Ojha P, Wiggs JL, Pasquale LR. The genetics of intraocular pressure. Semin Ophthalmol. 2013;28(5-6):301–305. doi:10.3109/08820538.2013.825291

52. Wang Z, Wiggs JL, Aung T, Khawaja AP, Khor CC. The genetic basis for adult onset glaucoma: Recent advances and future directions. Prog Retin Eye Res. 2022;90:101066. doi:10.1016/j.preteyeres.2022.101066

53. Khawaja AP, Cooke Bailey JN, Wareham NJ, et al. Genome-wide analyses identify 68 new loci associated with intraocular pressure and improve risk prediction for primary open-angle glaucoma. Nat Genet. 2018;50(6):778–782. doi:10.1038/s41588-018-0126-8

54. Liu W, Liu Y, Qin XJ, Schmidt S, Hauser MA, Allingham RR. AQP1 and SLC4A10 as candidate genes for primary open-angle glaucoma. Mol Vis. 2010;16:93–97.

55. Jian J, Bao X, Wang J, et al. An Integrated muti-omics cell atlas of the human trabecular meshwork and ciliary body. bioRxiv. Preprint posted online June 22, 2026:2026.06.17.732980. doi:10.64898/2026.06.17.732980

56. Kizhatil K, Simón M, Balasubramanian R, John SWM. Schlemm’s Canal Development and Implications for Glaucoma Treatment. Annu Rev Vis Sci. Published online May 6, 2026. doi:10.1146/annurev-vision-121423-013134

57. Perkins LA, Anderson CJ, Novelli EM. Targeting P-Selectin Adhesion Molecule in Molecular Imaging: P-Selectin Expression as a Valuable Imaging Biomarker of Inflammation in Cardiovascular Disease. J Nucl Med. 2019;60(12):1691–1697. doi:10.2967/jnumed.118.225169

58. He W, Qiu H, Feng Y, et al. P-selectin overexpression impairs hematopoietic stem cell homeostasis via inflammatory receptor-mediated proliferation and differentiation. Cell Death Dis. 2025;16(1):745. doi:10.1038/s41419-025-08050-9

59. Sasaki Y, Ohsawa K, Kanazawa H, Kohsaka S, Imai Y. Iba1 is an actin-cross-linking protein in macrophages/microglia. Biochem Biophys Res Commun. 2001;286(2):292–297. doi:10.1006/bbrc.2001.5388

60. Shiose Y, Kawase Y. A new approach to stratified normal intraocular pressure in a general population. Am J Ophthalmol. 1986;101(6):714–721. doi:10.1016/0002-9394(86)90776-2

61. Ajoolabady A, Pratico D, Vinciguerra M, Lip GYH, Franceschi C, Ren J. Inflammaging: mechanisms and role in the cardiac and vasculature. Trends Endocrinol Metab. 2023;34(6):373–387. doi:10.1016/j.tem.2023.03.005

62. Nguyen TQT, Cho KA. Targeting immunosenescence and inflammaging: advancing longevity research. Exp Mol Med. 2025;57(9):1881–1892. doi:10.1038/s12276-025-01527-9

63. Hearps AC, Martin GE, Angelovich TA, et al. Aging is associated with chronic innate immune activation and dysregulation of monocyte phenotype and function. Aging Cell. 2012;11(5):867–875. doi:10.1111/j.1474-9726.2012.00851.x

64. Solana R, Pawelec G, Tarazona R. Aging and innate immunity. Immunity. 2006;24(5):491–494. doi:10.1016/j.immuni.2006.05.003

65. Liu KC, Grimsrud AO, Suarez MF, et al. Resident tissue macrophages maintain intraocular pressure homeostasis. Immunity. 2026;59(4):943–952.e4. doi:10.1016/j.immuni.2026.01.025

66. Yan J, Chen S, Yi Z, et al. The role of p21 in cellular senescence and aging-related diseases. Mol Cells. 2024;47(11):100113. doi:10.1016/j.mocell.2024.100113

67. Deng T, Yan G, Song X, et al. Deubiquitylation and stabilization of p21 by USP11 is critical for cell-cycle progression and DNA damage responses. Proc Natl Acad Sci U S A. 2018;115(18):4678–4683. doi:10.1073/pnas.1714938115

68. Wong CA, Read AT, Li G, et al. Segmental Outflow and Trabecular Meshwork Stiffness in an Ocular Hypertensive Mouse Model. Invest Ophthalmol Vis Sci. 2026;67(6):51. doi:10.1167/iovs.67.6.51

69. Karimi A, Aga M, Stanik A, et al. Matrix stiffness regulates traction forces, cytoskeletal dynamics, and collagen reorganization in trabecular meshwork cells in glaucoma. Matter. 2025;8(6):102094. doi:10.1016/j.matt.2025.102094

70. Overby DR, Ethier CR, Miao C, Kelly RA, Reina-Torres E, Stamer WD. The Factors Affecting the Stability of IOP Homeostasis. Invest Ophthalmol Vis Sci. 2024;65(6):4. doi:10.1167/iovs.65.6.4

71. Li H, Kuhn M, Kelly RA, et al. Targeting YAP/TAZ mechanosignaling to ameliorate stiffness-induced Schlemm’s canal cell pathobiology. Am J Physiol Cell Physiol. 2024;326(2):C513–C528. doi:10.1152/ajpcell.00438.2023

72. Yemanyi F, Vranka J, Raghunathan VK. Crosslinked Extracellular Matrix Stiffens Human Trabecular Meshwork Cells Via Dysregulating β-catenin and YAP/TAZ Signaling Pathways. Invest Ophthalmol Vis Sci. 2020;61(10):41. doi:10.1167/iovs.61.10.41

73. Ozel AB, Moroi SE, Reed DM, et al. Genome-wide association study and meta-analysis of intraocular pressure. Hum Genet. 2014;133(1):41–57. doi:10.1007/s00439-013-1349-5

74. Choquet H, Thai KK, Yin J, et al. A large multi-ethnic genome-wide association study identifies novel genetic loci for intraocular pressure. Nat Commun. 2017;8:2108. doi:10.1038/s41467-017-01913-6

75. Gao XR, Huang H, Nannini DR, Fan F, Kim H. Genome-wide association analyses identify new loci influencing intraocular pressure. Hum Mol Genet. 2018;27(12):2205–2213. doi:10.1093/hmg/ddy111

76. Kizhatil K, Ryan M, Marchant JK, Henrich S, John SWM. Schlemm’s canal is a unique vessel with a combination of blood vascular and lymphatic phenotypes that forms by a novel developmental process. PLoS Biol. 2014;12(7):e1001912. doi:10.1371/journal.pbio.1001912

77. Thomson BR, Souma T, Tompson SW, et al. Angiopoietin-1 is required for Schlemm’s canal development in mice and humans. J Clin Invest. 2017;127(12):4421–4436. doi:10.1172/JCI95545

78. Aspelund A, Tammela T, Antila S, et al. The Schlemm’s canal is a VEGF-C/VEGFR-3-responsive lymphatic-like vessel. J Clin Invest. 2014;124(9):3975–3986. doi:10.1172/JCI75395

79. Kim J, Park DY, Bae H, et al. Impaired angiopoietin/Tie2 signaling compromises Schlemm’s canal integrity and induces glaucoma. J Clin Invest. 2017;127(10):3877–3896. doi:10.1172/JCI94668

80. Wang B, Han J, Elisseeff JH, Demaria M. The senescence-associated secretory phenotype and its physiological and pathological implications. Nat Rev Mol Cell Biol. 2024;25(12):958–978. doi:10.1038/s41580-024-00727-x

81. Das H, Kumar A, Lin Z, et al. Kruppel-like factor 2 (KLF2) regulates proinflammatory activation of monocytes. Proc Natl Acad Sci U S A. 2006;103(17):6653–6658. doi:10.1073/pnas.0508235103

82. Guixé-Muntet S, de Mesquita FC, Vila S, et al. Cross-talk between autophagy and KLF2 determines endothelial cell phenotype and microvascular function in acute liver injury. J Hepatol. 2017;66(1):86–94. doi:10.1016/j.jhep.2016.07.051

83. Dekker RJ, van Thienen JV, Rohlena J, et al. Endothelial KLF2 links local arterial shear stress levels to the expression of vascular tone-regulating genes. Am J Pathol. 2005;167(2):609–618. doi:10.1016/S0002-9440(10)63002-7

84. Linton MF, Yancey PG, Davies SS, et al. The Role of Lipids and Lipoproteins in Atherosclerosis. In: Feingold KR, Adler RA, Ahmed SF, et al., eds. Endotext. MDText.com, Inc.; 2000. Accessed July 6, 2026. http://www.ncbi.nlm.nih.gov/books/NBK343489/

85. Sangwung P, Zhou G, Nayak L, et al. KLF2 and KLF4 control endothelial identity and vascular integrity. JCI Insight. 2017;2(4):e91700. doi:10.1172/jci.insight.91700

86. Atkins GB, Wang Y, Mahabeleshwar GH, et al. Hemizygous Deficiency of Kruppel-like Factor 2 Augments Experimental Atherosclerosis. Circ Res. 2008;103(7):690–693. doi:10.1161/CIRCRESAHA.108.184663

87. Gupta GS. Selectins and Associated Adhesion Proteins in Inflammatory disorders. Animal Lectins: Form, Function and Clinical Applications. Published online March 20, 2012:991–1026. doi:10.1007/978-3-7091-1065-2_44

88. Tinoco R, Otero DC, Takahashi AA, Bradley LM. PSGL-1: A New Player in the Immune Checkpoint Landscape. Trends Immunol. 2017;38(5):323–335. doi:10.1016/j.it.2017.02.002

89. Zhong C, Wang L, Liu Y, et al. PSGL-1 is a phagocytosis checkpoint that enables tumor escape from macrophage clearance. Sci Immunol. 2025;10(108):eadn4302. doi:10.1126/sciimmunol.adn4302

90. Weyrich AS, Elstad MR, McEver RP, et al. Activated platelets signal chemokine synthesis by human monocytes. J Clin Invest. 1996;97(6):1525–1534. doi:10.1172/JCI118575

91. Klein BE, Klein R, Sponsel WE, et al. Prevalence of glaucoma. The Beaver Dam Eye Study. Ophthalmology. 1992;99(10):1499–1504. doi:10.1016/s0161-6420(92)31774-9

92. Hao Y, Hao S, Andersen-Nissen E, et al. Integrated analysis of multimodal single-cell data. Cell. 2021;184(13):3573–3587.e29. doi:10.1016/j.cell.2021.04.048

93. Young MD, Behjati S. SoupX removes ambient RNA contamination from droplet-based single-cell RNA sequencing data. Gigascience. 2020;9(12):giaa151. doi:10.1093/gigascience/giaa151

94. van Zyl T, Yan W, McAdams AM, Monavarfeshani A, Hageman GS, Sanes JR. Cell atlas of the human ocular anterior segment: Tissue-specific and shared cell types. Proceedings of the National Academy of Sciences. 2022;119(29):e2200914119. doi:10.1073/pnas.2200914119

95. Timmons JA, Szkop KJ, Gallagher IJ. Multiple sources of bias confound functional enrichment analysis of global -omics data. Genome Biol. 2015;16(1):186. doi:10.1186/s13059-015-0761-7

96. Yu G, Wang LG, Han Y, He QY. clusterProfiler: an R Package for Comparing Biological Themes Among Gene Clusters. OMICS: A Journal of Integrative Biology. 2012;16(5):284–287. doi:10.1089/omi.2011.0118

97. Xin Y, Lyu P, Jiang J, et al. LRLoop: a method to predict feedback loops in cell–cell communication. Bioinformatics. 2022;38(17):4117–4126. doi:10.1093/bioinformatics/btac447

98. Browaeys R, Saelens W, Saeys Y. NicheNet: modeling intercellular communication by linking ligands to target genes. Nat Methods. 2020;17(2):159–162. doi:10.1038/s41592-019-0667-5

